# Genetic basis of historical pea mutants that hyper-accumulate iron

**DOI:** 10.1101/2023.06.05.543728

**Authors:** Sophie A. Harrington, Marina Franceschetti, Janneke Balk

**Affiliations:** Department of Biochemistry and Metabolism, John Innes Centre, Norwich NR4 7UH, UK

**Keywords:** iron, hemerythrin, RNA-seq, *Pisum sativum*, *Medicago truncatula*

## Abstract

The *Pisum sativum* (pea) mutants *degenerate leaves* (*dgl*) and *bronze* (*brz*) accumulate large amounts of iron in leaves. First described several decades ago, the two mutants have provided important insights into iron homeostasis in plants but the underlying mutations have remained unknown. Using exome sequencing we identified an in-frame deletion associated with *dgl* in a *BRUTUS* homologue. The deletion is absent from wild type and the original parent line. BRUTUS belongs to a small family of E3 ubiquitin ligases acting as negative regulators of iron uptake in plants. The *brz* mutation was previously mapped to chromosome 4, and superimposing this region to the pea genome sequence uncovered a mutation in *OPT3*, encoding an oligopeptide transporter with a plant-specific role in metal transport. The causal nature of the mutations was confirmed by additional genetic analyses. Identification of the mutated genes rationalises many of the previously described phenotypes and provides new insights into shoot-to-root signalling of iron deficiency. Furthermore, the non-lethal mutations in these essential genes suggest new strategies for biofortification of crops with iron.

**Significance statement:** Two iron-accumulating pea mutants first described more than 30 years ago have greatly contributed to our understanding of iron homeostasis in plants, but the mutations were never identified. Here we show that the phenotypes are caused by mutations in the *BRUTUS* and *OPT3* genes and how this leads to specific defects in iron signalling and leaf development.

## Introduction

Prior to the rise of Arabidopsis as a plant model organism, genetic studies in pea contributed greatly to our understanding of the mechanisms of inheritance, and also helped identify several genetic loci for morphological and nutritional traits in plants (Ellis *et al*, 2011). As part of these studies, two iron-accumulating pea mutants were independently isolated: the *dgl* mutant, from an X-ray mutagenized population (Gottschalk, 1987), and the *brz* mutant, also named E107, generated using ethyl methanesulfonate (EMS) (Kneen *et al*, 1990). In addition to a root nodulation phenotype, the leaves of both mutants develop striking bronze spots and senesce prematurely (**Fig. 1A**, bottom panel). The spots are necrotic tissue caused by toxic levels of iron, which accumulates 10 to 100-fold in older leaves (Welch & LaRue, 1990; **Fig. 1B, C**). Despite the similar phenotypes, genetic analysis showed that the *brz* and *dgl* loci segregate independently (Kneen *et al*, 1990). The *brz* mutation was mapped to a large segment of chromosome 4, but was not further fine-mapped or cloned. For *dgl*, difficulties with distinguishing heterozygotes within the F2 population due to variability in iron concentration thwarted efforts to obtain any idea of its genome location (Kneen *et al*, 1990).

**Figure 1.**
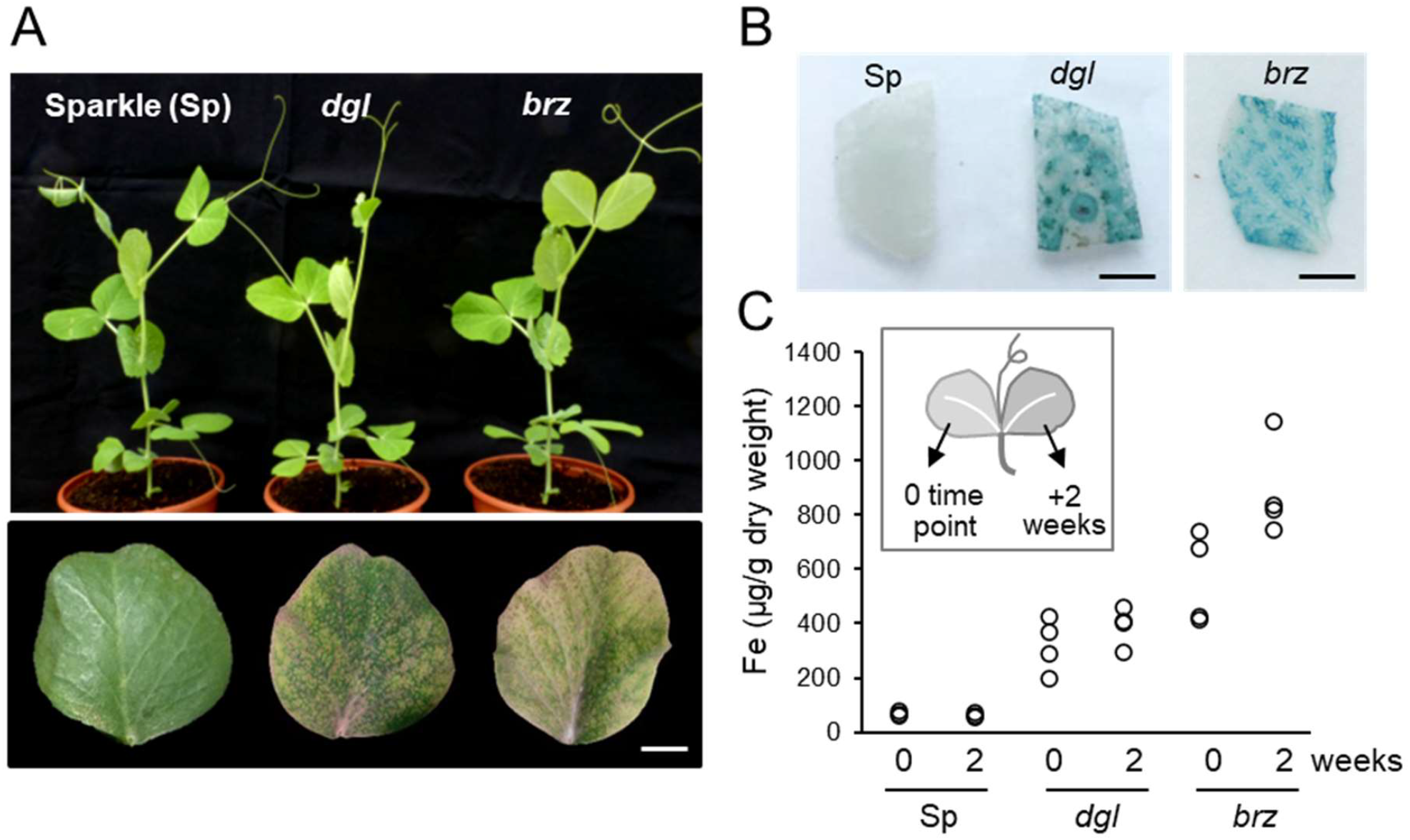
The *dgl* and *brz* mutants in pea (*Pisum sativum* L.) hyper-accumulate iron in leaves. A. Top panel: Three-week old plants of the pea variety Sparkle (Sp) and the mutants *dgl* and *brz* in the same genetic background. Bottom panel: Lower leaflet of 5-week-old plants, showing yellowing and brown spots. Scale bar = 1 cm. B. Iron staining (blue) of leaf pieces from 5-week-old plants. Scale bar = 2 mm. C. Iron concentrations in leaflets of the same leaf, measured two weeks apart using a colorimetric assay. Dots represent measurements of individual plants.

Despite not knowing the mutated genes, seminal physiological studies of the *dgl* and *brz* mutants demonstrated the existence of a Fe^3+^ reductase activity induced by iron deficiency (Welch & LaRue, 1990; Grusak *et al*, 1990) well before the corresponding gene was cloned from Arabidopsis (Robinson *et al*, 1999). In addition, grafting wild-type shoots onto mutant roots and vice versa revealed the existence of a shoot-to-root signal for iron deficiency (Welch & LaRue, 1990; Grusak & Pezeshgi, 1996; García *et al*, 2013), the molecular nature of which is still unknown.

Because iron is toxic and controlled by tight homeostasis mechanisms, there is limited genetic variation for the iron concentration in crops and mutant screens rarely turn up iron-accumulating mutants (Connorton & Balk, 2019; Lahner *et al*, 2003). Therefore, identifying the *dgl* and *brz* mutations could be important both for our general understanding of iron homeostasis and to help design strategies for biofortifying crops.

Finding genetic loci is greatly facilitated by a genome sequence, but owing to its large size (∼4.45 Gb), a draft of the pea genome was not published until recently (Kreplak *et al*, 2019). This confirmed the close relationship and extensive co-localization of genetic loci (synteny) with the *Medicago truncatula* genome. Making use of this new resource, we show that *dgl* is caused by a short deletion in *BRUTUS*, a putative iron sensor and negative regulator of iron uptake, and that *brz* is associated with mutations in the iron transporter *OPT3* in both pea and Medicago.

## Results and Discussion

### *dgl* is associated with a small in-frame deletion in *BRUTUS*

To investigate the transcriptional basis of iron accumulation in the *dgl* pea mutant and, if possible, to identify the causative mutation, we carried out RNA sequencing on leaves of *dgl* and the wild-type cultivar Sparkle, into which the original *dgl* mutation was introgressed by at least five backcrosses (Marentes & Grusak, 1998). We found 86 differentially expressed genes which were highly enriched for genes involved in iron homeostasis (**Suppl. Table 1 and 2**). These included four different ferritin genes with massively induced expression, in agreement with previous reports of increased ferritin protein in *dgl* leaves and seeds by electron microscopy, Western blot analysis and iron-stained native gels (Becker *et al*, 1998; Marentes & Grusak, 1998). Also upregulated were vacuolar iron transporters and a gene involved in zinc detoxification (PCR2), whereas various sugar metabolism genes are downregulated (**Suppl. Table 2**).

Alignment of the RNA reads to the pea genome sequence (Kreplak *et al*, 2019) followed by identification of SNPs and Indels in *dgl* relative to Sparkle confirmed that the lines were near-isogenic (**Suppl. Fig. 1**). Because the mutant was generated by X-rays, we focussed on the Indels as candidate genetic polymorphisms (**Table 1**). Molecular marker analysis showed that 3 Indels located on chromosome 6 were not linked to the mutant phenotype (data not shown). However, a deletion located on chromosome 1 co-segregated invariably with the high-iron phenotype in the F2 population (n = 11 mutants out of 44 plants, **Suppl. Fig. 2**) and is not present in the original Dippes Gelbe Viktoria pea variety used for mutagenesis (**Fig. 2A**). The 15-bp deletion is located in exon 2 of *Psat1g036240*, corresponding to an in-frame deletion of five amino acids (**Fig. 2B**). A blastp search against the Arabidopsis proteome revealed that Psat1g036240 shares 66.4% identity with the BRUTUS (BTS) protein from Arabidopsis, an E3 ubiquitin ligase that acts as a negative regulator of iron uptake (Selote *et al*, 2015; Rodríguez-Celma *et al*, 2019). The five deleted amino acids, QTSLS, are located in the first hemerythrin motif (**Fig. 2B**) and are semi-conserved in BTS sequences across the green lineage. Loss of these residues is predicted to displace two of the seven amino acid ligands that coordinate a diiron centre, thus potentially weakening iron binding and affecting the proposed oxygen and/or iron sensing function of this domain (**Fig. 2C; Suppl. Fig. 3**).

**Table 1.** Indels in *dgl* identified by exome mapping. Four indels were identified as candidate X-ray mutation for *dgl* by comparing RNA-seq data from *dgl* and wild-type variety Sparkle mapped to the pea genome (variety Cameor). Only the deletion on chromosome 1 co-segregated with the *dgl* phenotype (see Suppl. Fig. 2).

| Chromosome / Linkage Group | Indel size | Gene | Outcome of mutation | Arabidopsis BLAST hits | Co-segregates? |
| --- | --- | --- | --- | --- | --- |
| Chr1 / LG6 | 15 bp | *Psat1g036240* | In-frame deletion of TSLSQ | BRUTUS | Yes |
| Chr6 / LG2 | 24 bp | *Psat6g181240* | In-frame deletion of SMQQPLPS | Decapping 5 (DCP5) | No |
| Chr6 / LG2 | 15 bp | *Psat6g182800* | In-frame deletion of QTLPL | Mediator associated protein 1-like | No |
| Chr6 / LG2 | 21 bp | *Psat6g183600* | Insertion causes premature stop, deleting final VLVQ | No hits | No |

**Figure 2.**
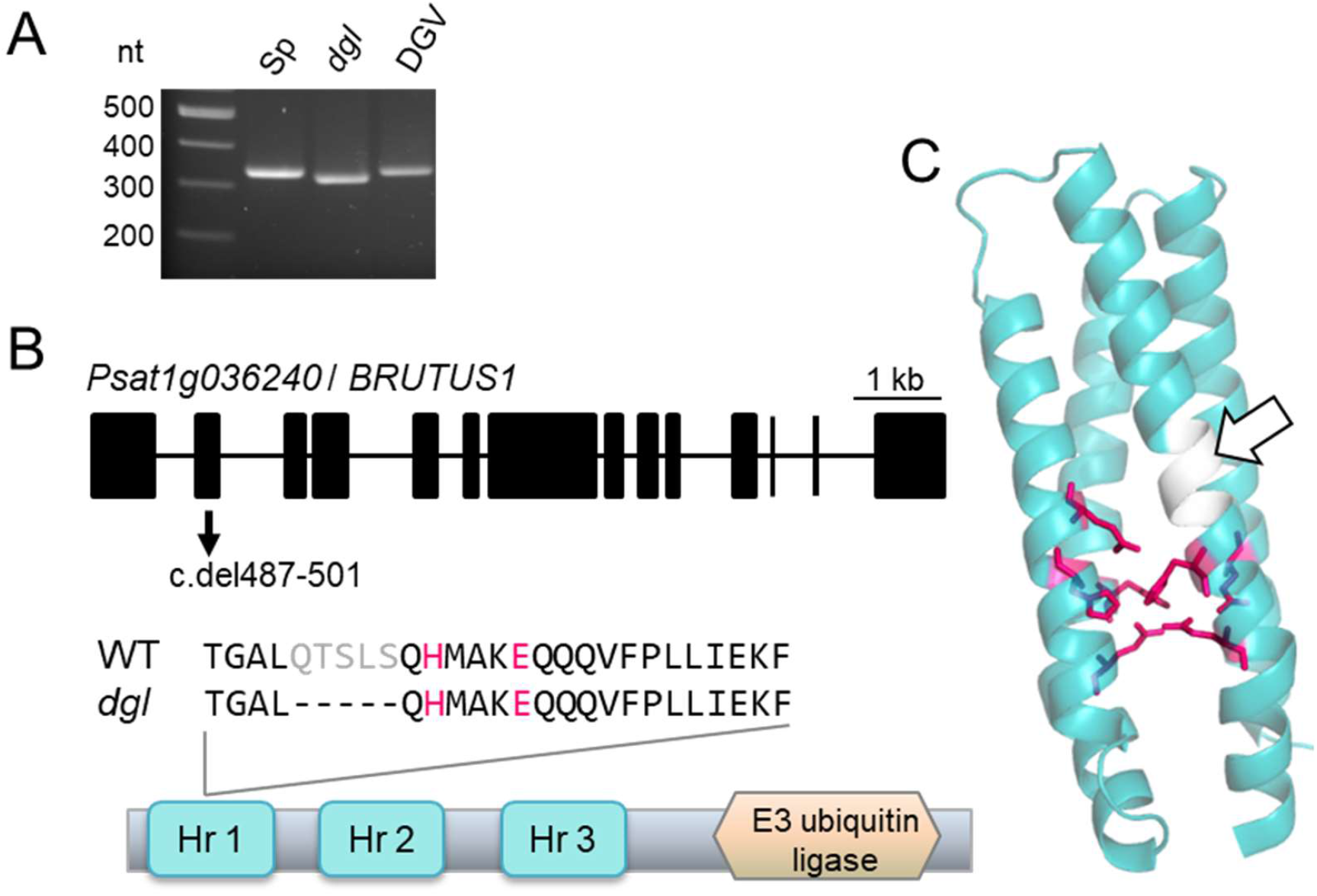
*dgl* is associated with a small in-frame deletion in *BRUTUS*. A. PCR analysis with primers spanning the 15-bp deletion associated with *dgl* show that the deletion is absent from the near-isogenic wild-type Sparkle (Sp) variety and also from the Dippes Gelbe Viktoria (DGV) variety that was originally used for mutagenesis (Gottschalk, 1987). B. The *dgl* mutant has a mutation in exon 2 of the gene *Psat1g036240*, encoding a BRUTUS homolog. The 15-bp in-frame deletion removes five amino acids (QTSLS) of the first hemerythrin domain, close to two iron-binding residues (H and E) in a conserved HxxxE motif (in red). C. Protein model of the Hr1 domain, with ligands binding the di-iron centre in magenta. The deleted 5 amino acids in the *dgl* mutant are rendered in white and marked with a white arrow. The deletion is predicted to displace two of the seven iron-binding ligands, see Suppl. Fig. 3.

### *brz* is associated with a point mutation in *OPT3*

The *brz* mutation was previously mapped to the tip of chromosome 4 between the phenotypic markers *lat* (latum, Latin for wide leaves) and *was* (waxy stipules) (Kneen *et al*, 1990; Ellis and Poyser, 2002; **Fig. 3A**). These two loci have not been identified, but closely linked genes that have been cloned suggested that *brz* lies between *Psat4g001240* and *Psat4g005920*, an interval of more than 450 genes. Near the middle of this interval are the likely gene candidates *Psat4g003000* and *Psat4g003080*, paralogous genes encoding the oligopeptide transporter OPT3. This member of the large OPT transporter family is unique to vascular plants and forms a separate phylogenetic clade with a single origin. Mutant studies in Arabidopsis showed that OPT3 has an essential function in transporting iron from the xylem to the phloem (Zhai *et al*, 2014; Mendoza-Cózatl *et al*, 2014) and also mediates copper transport (Chia *et al*, 2023). Publicly available transcriptome data of pea suggests that *Psat4g003000* does not have meaningful expression levels whereas *Psat4g003080* is expressed in all plant organs, particularly in young leaves and stems (**Suppl. Fig. 4B**). Sequencing of the coding sequence identified a C > T mutation, consistent with the effect of EMS as a mutagen, that co-segregated with the iron-accumulating phenotype (**Suppl. Fig. 5**). The missense mutation changes a conserved leucine residue into phenylalanine (Leu466Phe, **Fig. 3B**). Protein modelling suggests that Leu466 is part of an alpha helix oriented inwards (**Fig. 3C**) and substitution to a larger, more hydrophobic side group may interfere with the transport mechanism.

**Figure 3.**
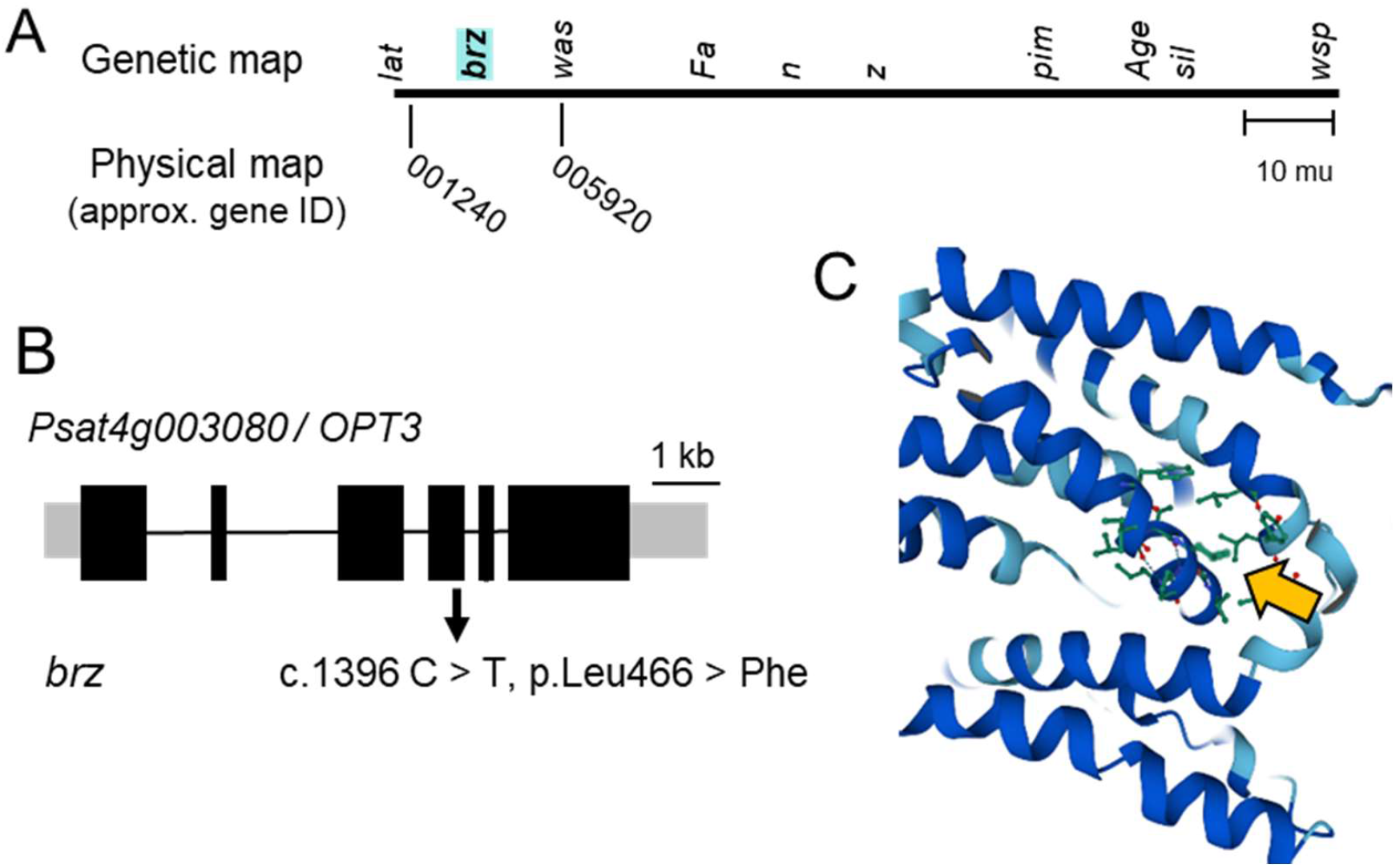
*brz* is associated with a point mutation in *OPT3*. A. The *brz* mutation was previously mapped to the tip of chromosome 4, between the phenotypic loci *lat* and *was* (Kneen et al., 1990; Ellis & Poyser, 2002). Neigbouring loci that have been cloned were used to obtain a physical interval between genes *Psat4g001240* and *Psat4g005920*. B. The *brz* mutant contains a C to T mutation in exon 4 of the gene *Psat4g003080*, encoding the oligopeptide transporter OPT3, resulting in substitution of the highly conserved leucine 466 to phenylalanine. C. Protein model of Arabidopsis OPT3, centred on Leu461 (equivalent to Leu466 in pea OPT3, yellow arrow).

### Further genetic evidence for the mutations of *dgl* and *brz*

To confirm that the identified mutations are causally linked to the *dgl* and *brz* phenotypes, we used different approaches. First, genetic complementation in pea was attempted by transiently expressing the wild-type pea *BTS1* cDNA in *dgl* seedlings, using *Agrobacterium*-mediated leaf infiltration. This did not decrease or prevent iron accumulation in the infiltrated leaves, either because expression was too low (GFP levels expressed using a separate construct were also low) or because iron accumulation in this mutant is controlled systemically and cannot be suppressed locally. Stable transformation of pea has been reported but is technically challenging. Therefore, as a second approach, we applied TILLING (Targeting Induced Local Lesions IN Genomes) to the syntenic Medicago genes. An EMS-mutagenized population was screened for sequence polymorphisms in a 1.5 - 2 kb DNA region overlapping with or close to, respectively, the position of the *brz* and *dgl* mutations.

For *OPT3*, we isolated four non-synomymous mutations altering conserved amino acid residues. Of those, Medicago plants with Pro529Leu or Pro618Leu presented with bronze spots on the leaves and intense iron staining (**Fig. 4A**). Pro618Leu homozygous seedlings segregated at a low frequency and had a severe growth phenotype, confirming the critical function of *OPT3* in Medicago. For *BTS1*, an unusually small number of mutations was found in the selected region (exons 7 and 8, corresponding to the Hr 3 domain, Fig. 2C). Of the five non-synomymous mutations, only two affected conserved amino acids. Asp912Asn had no discernable phenotype, whereas Asp730Asn homozygous plants could not be found among the offspring of a heterozygous plant. This suggests that *BTS1* is an essential gene in Medicago, similar to *BTS* being essential for embryo development in Arabidopsis (Selote *et al*, 2015), and prevented further studies in young seedlings or mature plants.

**Figure 4.**
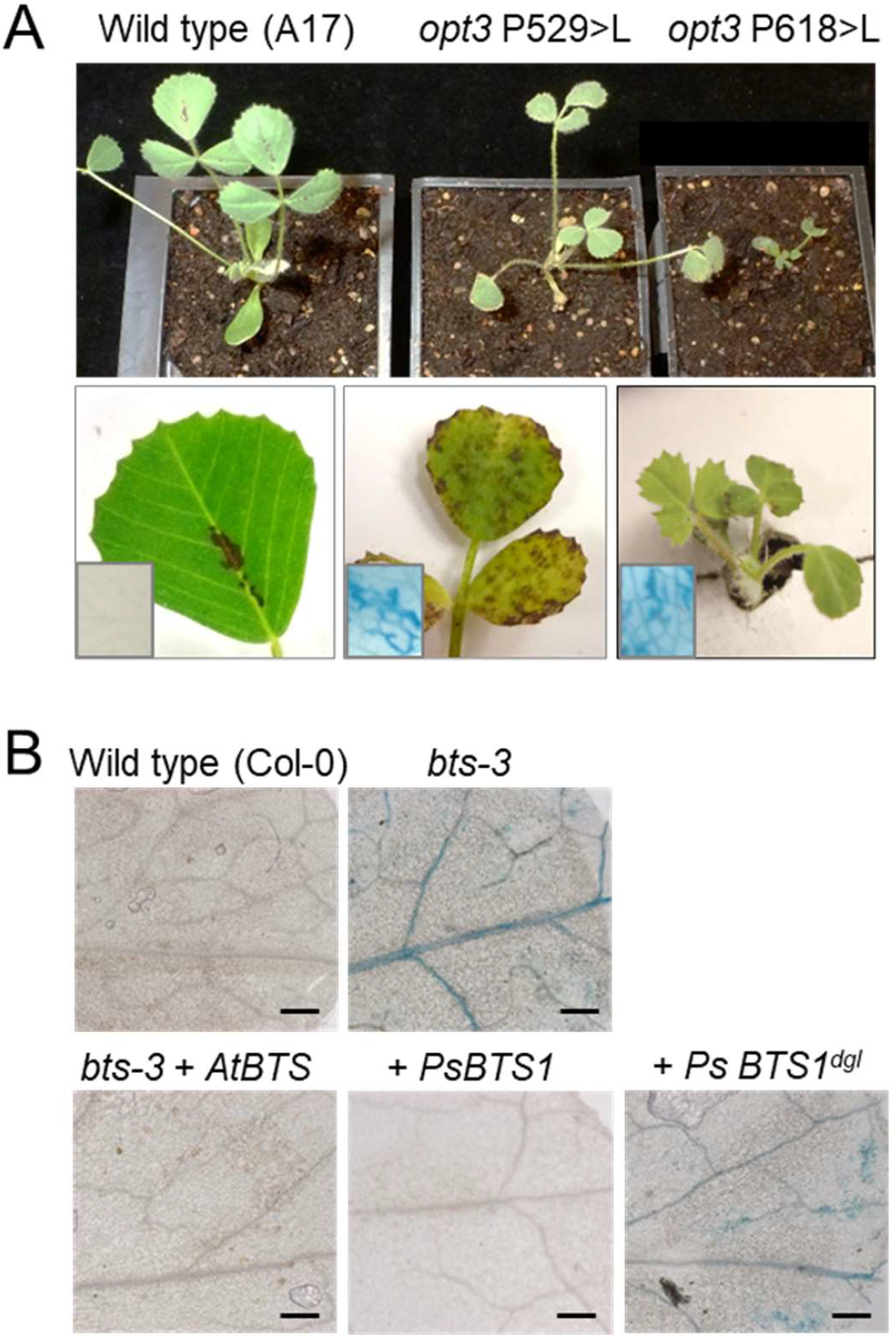
Further genetic evidence for the mutated genes in *dgl* and *brz*. A. Mutations in *Medicago truncatula OPT3*, which is syntenic with pea *OPT3*/*Psat4g003080*, phenocopy the pea *brz* mutant, including necrotic leaf spots and iron accumulation. Wild type (A17) and mutants in *OPT3* (*Medtr6g083900*), c.C1586>T (p. P529L) and c.C1853>T (p. P618L) grown on soil (top); close up of the leaves (middle); and leaf sectors stained for iron (insets). The dark purple spots in the middle of wild-type leaves are anthocyanin. Scale bars in the top panel are 1 cm. B. The wild-type pea (*Ps*) *BTS1* coding sequence genetically complements the iron accumulation phenotype of the Arabidopsis *bts-3* mutant, but the pea *BTS1*-*dgl* sequence does not. Detail of rosette leaves stained for iron, imaged by light microscopy. Scale bars are 0.2 mm. Genotyping data of the plants can be found in Suppl. Fig. 6B.

**Figure 5.**
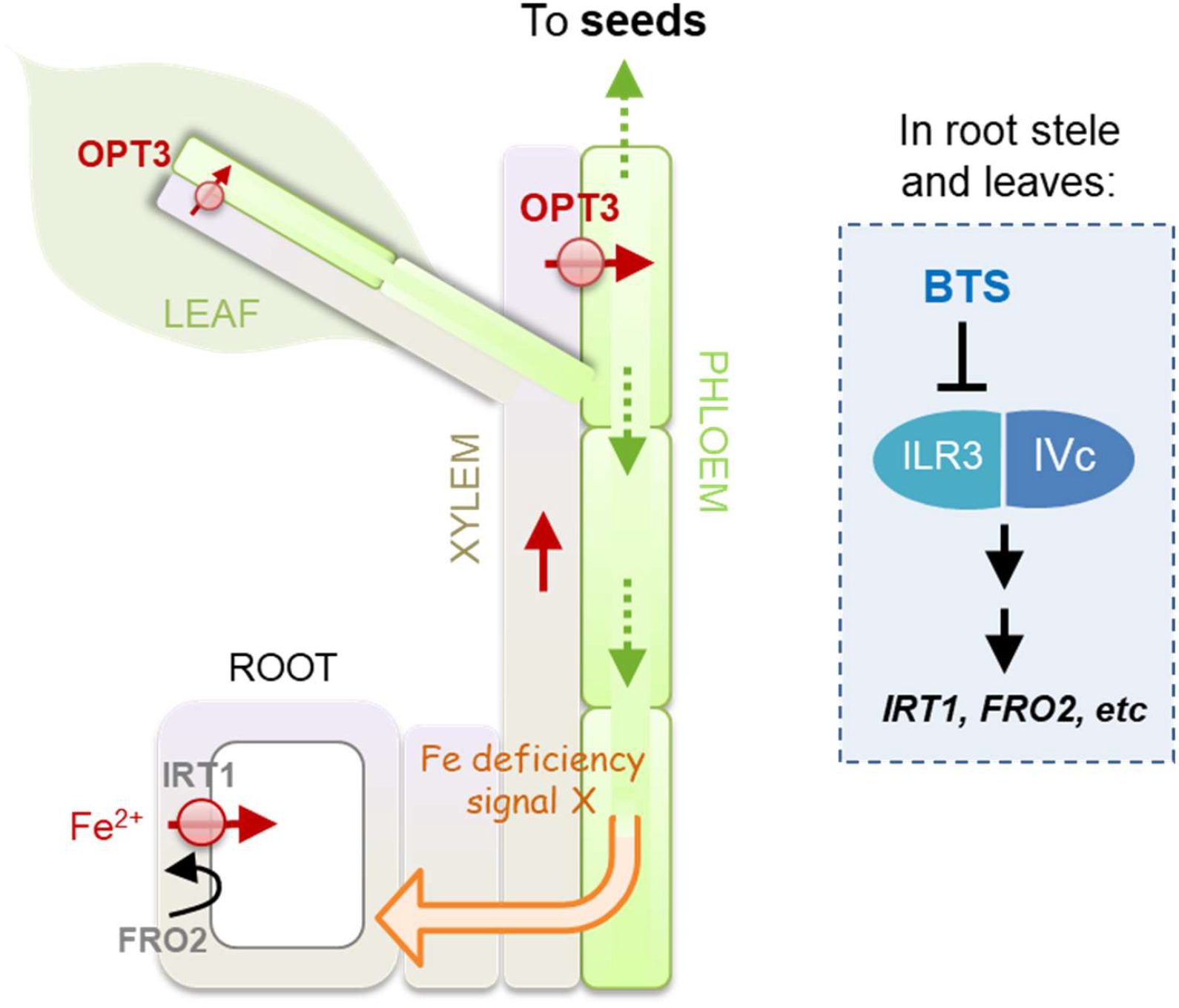
Proposed mechanism of iron accumulation in *dgl* (*bts1*) and *brz* (*opt3*) mutants. Reduced function of the E3 ubiquitin ligase BRUTUS (BTS) or the oligopeptide transporter OPT3 in the *dgl* and *brz* mutants, respectively, leads to constitutive activity of the iron uptake pathway in roots (Welch & LaRue, 1990; Grusak *et al*, 1990). The pea *BTS1* gene is expressed predominantly in shoots (https://urgi.versailles.inra.fr/), in agreement with Arabidopsis *BTS* promoter activity and high transcript levels in leaves, root stele and embryo cotyledons (Selote *et al*, 2016; Hindt *et al*, 2017). Arabidopsis BTS interacts with the transcription factor ILR3 to target it for degradation (Selote *et al*, 2016), although evidence of ubiquitination is currently lacking. Decreased activity of BTS leads to increased levels (and transcriptional activity) of ILR3, which indirectly leads to enhanced transcription of iron uptake genes. Iron accumulates in minor and major veins and also in seeds. AtOPT3 is expressed in the phloem (Zhai et al, 2014; Mendoza-Cózatl et al, 2014) and mediates transport of iron and copper from the xylem into the phloem. Impairment of OPT3 function leads to accumulation of iron in the xylem, in particular minor veins, and to decreased iron levels in the phloem and in seeds. Iron deficiency in the phloem triggers a signal, the nature of which is still unknown, that induces the expression of iron uptake genes in the roots and enhanced activity of the ferric reductase FRO2.

Because no other exome sequence polymorphisms were found in close range of the 15-bp deletion associated with *dgl*, it is rather unlikely that a different, yet genetically linked, mutation causes the iron-accumulating phenotype. Nevertheless, as a third approach, we tested for complementation of the Arabidopsis *bts-3* mutant (Hindt *et al*, 2017) with the coding sequences of wild-type pea *BTS1* or the *dgl*-associated mutant gene. As a control, plants were transformed with Arabidopsis *BTS*. A previous study showed that *bts-3* mutant plants, which carry a point mutation in the E3 ligase domain, accumulated significant amounts of iron and had a severe growth defect (Hindt *et al*, 2017). We found that in the T2 generation growth was variable, perhaps because of epigenetic effects. However, iron accumulation visualized by Perls’ staining, which was most prominent in the veins, was clearly suppressed in *bts-3* plants transformed with *AtBTS* or pea *BTS1*, but not with the *dgl*-associated variant of *BTS1* (**Fig. 4B; Suppl. Fig. 6**). These results demonstrate that pea *BTS1* is a functional orthologue of Arabidopsis *BTS* and that the 15-bp deletion is deleterious for BTS function.

### Phenotypic differences between *dgl* and *brz* are associated with the specific functions of BTS and OPT3, respectively, in iron homeostasis

The pea *dgl* and *brz* mutants have an unusual history in that detailed reports of their phenotypes have been published since 1990, which can only now be matched to the affected genes. While both mutants accumulate large amounts of iron in leaves, the pattern of accumulation differs in subtle ways. In the *brz* / *opt3* mutant, iron staining is restricted to the narrow ends of leaf veins, consistent with high expression of the Arabidopsis *OPT3* gene in the minor veins (Zhai *et al*, 2014) (**Suppl. Fig. 7**). By contrast, the *dgl* mutant accumulates iron in both minor and major veins, matching the promoter activity of Arabidopsis *BTS*, which can reasonably be extrapolated to the pea *BTS1* gene. *dgl* mutants also accumulate iron in the seeds, whereas *brz* seeds have less iron compared to wild type (Grusak, 1994; Marentes & Grusak, 1998). Seeds derive their iron stores from the phloem (**Fig. 5**), and iron loading into the phloem is strongly impaired in *opt3* mutants. Moreover, transcript levels of *OPT3* are very low in Arabidopsis seeds, whereas *BTS* is highly expressed in the embryo cotyledons (Selote *et al*, 2015; Zhai *et al*, 2014). From what we know of the function of BTS and its main ubiquitination target ILR3 (Selote *et al*, 2015; Akmakjian *et al*, 2021), BTS suppresses the activity of the transcriptional cascade for iron uptake and, via ILR-PYE interaction, regulates iron remobilization (**Fig. 5**). *dgl* seeds accumulate up to 4 times more iron, which is less than in leaves, probably reflecting a balance between limited iron supply through the phloem (with iron not being remobilized from leaf veins) and upregulated iron uptake in the embryo.

Grafting experiments demonstrated the existence of a shoot-to-root signal leading to constitutive iron uptake in the roots of *dgl* and *brz* mutants (Welch & LaRue, 1990; Grusak & Pezeshgi, 1996). The signal is thought to be generated by iron deficiency in the phloem, but its molecular nature remains to be identified. Interestingly, *brz* shoots also suppressed the number of symbiotic root nodules (Huynh & Guinel, 2020) and it is possible that the same signalling molecule leads to both physiological outputs. The role of neither BTS or OPT3 in iron homeostasis during nodule development has been properly investigated (Day & Smith, 2021). *opt3* mutants will be invaluable to shed light on the question of whether iron is delivered to nodules via the xylem or phloem, or both. *bts* mutants could help lift the lid on the regulation of iron homeostasis in nodules, including downstream transcription factors.

Increasing the mineral micronutrient of crops, or biofortification, is an important area of research to combat hidden hunger and to provide nutritious plant-based diets in the face of climate change. The identified mutations indicate that either BTS or OPT3 function could be modified to increase the iron content of vegetative tissues. Genetic variants of *BTS* could also enhance iron in seeds. However, there is a fine balance between increasing iron and toxicity symptoms, and null mutants of *BTS* and *OPT3* are embryo-lethal (Selote *et al*, 2015; Zhai *et al*, 2014; Mendoza-Cózatl *et al*, 2014). Interestingly, the pea *dgl* mutant has relatively minor growth defects, indicating that it is worth screening for weaker alleles, especially in the Hr1 domain. In summary, knowing the causal mutations in the historic *dgl* and *brz* mutants will help to further unravel the functional roles of these important iron homeostasis genes.

## Materials and Methods

### Plant material and growth

Seeds of *dgl* (JI3085) and the wild type ‘Sparkle’ (JI0427) were obtained from the John Innes Centre Germplasm Resource Unit (GRU), which were donated to the collection by Michael Grusak, then at Asgrow Seeds, Twin Falls, USA. Seeds of the EMS mutant E107, also named *brz* (JI2616) were also obtained from GRU, donated by Thomas LaRue, Cornell University, Ithaca, USA. The germination rate of *brz* seeds declines rapidly over time, and the mutant should be propagated every 3 - 4 years. The *dgl* mutant was originally generated in the pea variety Dippes Gelbe Viktoria (DGV, accession JI2413 in the GRU collection). Marentes & Grusak (1998) backcrossed *dgl* at least five times with the Sparkle variety, selected at the F3 stage for the high-iron phenotype. For segregation analysis, *dgl* and *brz* were crossed with line JI804 because of its contrasting phenotypic traits and suitable genetic markers. Plants were germinated on peat-based compost (Levington F2) and grown in a greenhouse with additional lighting in the winter and watering as required.

### Leaf iron staining and quantification

Leaf samples were stained for iron using Perls’ reagent as previously described (Meguro *et al*, 2007). Samples were mounted in 50% (v/v) glycerol and imaged on a Axio Zoom.V16 stereo microscope with an Axiocam 512 color camera (Zeiss). For measuring iron concentrations, dried leaf samples were digested in 0.25 ml nitric acid (69% w/v) and 0.25 ml hydrogen peroxide (30% w/v) at 90°C. After neutralizing with 1 ml ammonium acetate (15% w/v), samples were reduced with 0.1 ml ascorbic acid (4% w/v). Fe^2+^ was quantified using the colorimetric iron chelator ferene (3-(2-pyridyl)-5,6-bis-[2-(5-furyl-sulfonic acid)]-1,2,4-triazine, 0.1% w/v) and absorbance was measurement at 593 nm.

### RNA extraction and Illumina sequencing

Leaf tissue was sampled from three plants of each genotype (Sparkle and *dgl*) and snap-frozen in liquid N2. The frozen tissue was ground to a fine powder before RNA extraction using TRIzol® Reagent (ThermoFisher) and DNase treatment with TURBO DNase (ThermoFisher). The quality and quantity of RNA was verified with the Agilent Bioanalyzer RNA 6000 Nano assay before cDNA library preparation (250-300 bp insert) and Illumina Sequencing (PE 150, Novogene).

### Differential expression analysis

Illumina reads were pseudo-aligned against the *P. sativum* reference transcriptome (Kreplak *et al*, 2019) using Kallisto (Bray *et al*, 2016). Gene expression levels were determined using the R package Sleuth (Pimentel *et al*, 2017) using the Wald test, where we considered genes to be differentially expressed between genotypes with *q* < 0.05. Enrichment of GO terms was calculated using the R package goseq (Young *et al*, 2010).

### Identification of deletions

Illumina reads were aligned against the *P. sativum* reference genome (Kreplak *et al*, 2019) using the software *BWA-mem* (Li, 2013). The software transIndel (Yang R & Van Etten JL, 2018) was then used to identify the location of deletions and insertions using the default parameters except for DP = 1 (to capture all possible deletions, irrespective of gene expression level).

### PCR analysis to confirm allelic variation

DNA was extracted from individual F2 plants in 200 mM Tris-HCl pH 7.5, 250 mM NaCl, 25 mM EDTA and 0.5% (w/v) sodium dodecyl sulfate and precipitated with 50% (v/v) isopropanol. For *dgl*, PCR primers were designed to amplify across the region of *Psat1g036240* which contains the identified five codon deletion: GCGTGAAGAATGTAGCACAG and ACCTGCAATATTCAACCAGCA, see Suppl. Table 3. Amplified fragments were run on a 2% (w/v) agarose gel for 2 h at 70 V, allowing separation of the wild-type Sparkle band (334 bp) and the *dgl* band (319 bp). For *brz*, PCR primers were designed to span part of the *Psat4g003080* coding sequence: GACATATTGAGACAGAGCAGG and ATACCGAATCATGAACTGTGC, see Suppl. Table 3. The PCR product was purified and sequenced.

### Protein homology modelling

The first hemerythrin domain of pea BTS1 / Psat1g036240.1 (amino acids 55 – 184), the *dgl* variant of this domain, the full length OPT3 protein and the L466F variant were modelled using AlphaFold2 and visualised using PyMol version 2.5.2.

### Genetic complementation of Arabidopsis bts-3

Heterozygous *bts-3* plants (Hindt *et al*, 2017) were transformed with plasmid pICSL869550OD (SynBio) carrying either the Arabidopsis *BRUTUS* coding sequence (*BTS, AT3G18290*); the pea *BRUTUS1* coding sequence (*Psat1g036240*); or the pea *dgl* variant of *BRUTUS1* (lacking nucleotides 487-510). The coding sequences were placed downstream of the Arabidopsis *BRUTUS* promoter, nucleotides -1904 to -1, and upstream of the *ocs* terminator, using Golden Gate assembly. All constructs were verified by sequencing. T1 plants homozygous for *bts-3* were selected by PCR (see Suppl. Table 3 for primers) followed by restriction with PflMI, the recognition site of which is deleted in the *dgl* allele. Of these lines, T2 plants carrying the transgene were scored for growth and iron accumulation.

### Medicago truncatula TILLING

A M2 population of EMS-mutagenized *Medicago truncatula* was screened for genetic polymorphisms in *BTS1*/*Psat1g036240* and *OPT3*/*Psat4g003080*. For *BTS1*, primers MtBTS1-F1 and -R spanned exons 7 - 10 to maximize the ratio of exon:intron sequence; For *OPT3*, primers MtOPT3-F1 and -R3 spanned exons 4 - 6, surrounding the *brz* mutation. See Suppl. Table 3 for primer sequences. For the selected lines (Suppl. Table 4), seedlings were grown up and inspected for phenotypes and iron accumulation using Perls’ staining.

## Acknowledgements

We would like to thank the JIC Germplasm Resource Unit for providing pea seed stocks and Saleha Bakht for isolating Medicago TILLING mutants; JIC horticultural services for plant growth; Lucy Anderson and Elina Kondratovica for assistance with phenotyping; Jacob Pullin for generating the protein models; Michael Grusak, Noel Ellis and Julie Hofer for helpful discussions.

Funding for this project was provided by the Biotechnology and Biological Sciences Research Council grants BB/P012523/1, BB/T004363/1 and BB/V015095/1.

---

**Supplemental Figure 1.**
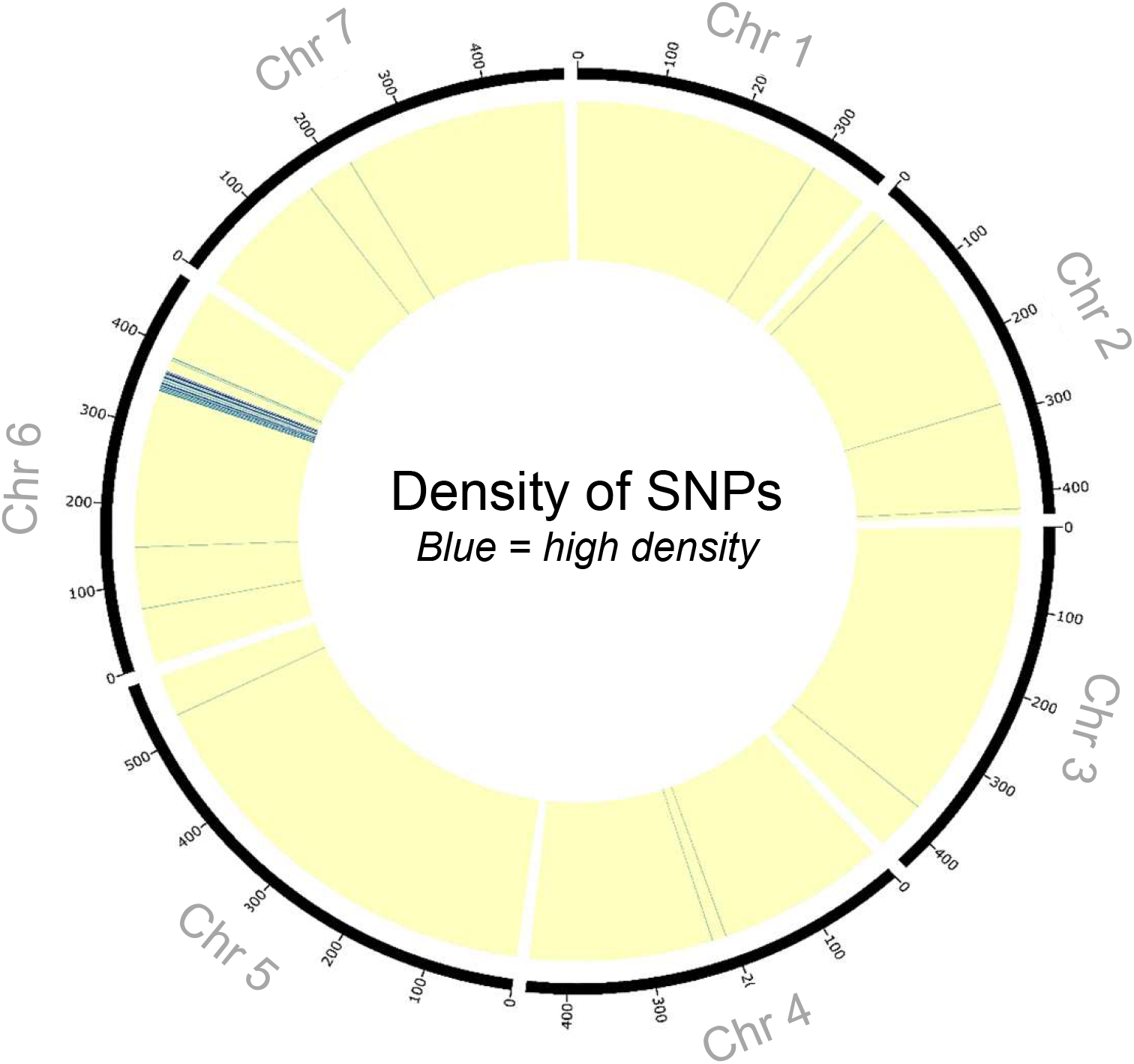
Exome mapping to identify the *dgl* mutation. Circular representation of the pea (*Pisum sativum* L.) genome to which the RNA-seq data from *dgl* and Sparkle (wild type) are mapped. Blue lines represent sequence polymorphisms and yellow indicates sequence identity between *dgl* and Sparkle.

**Supplemental Figure 2.**
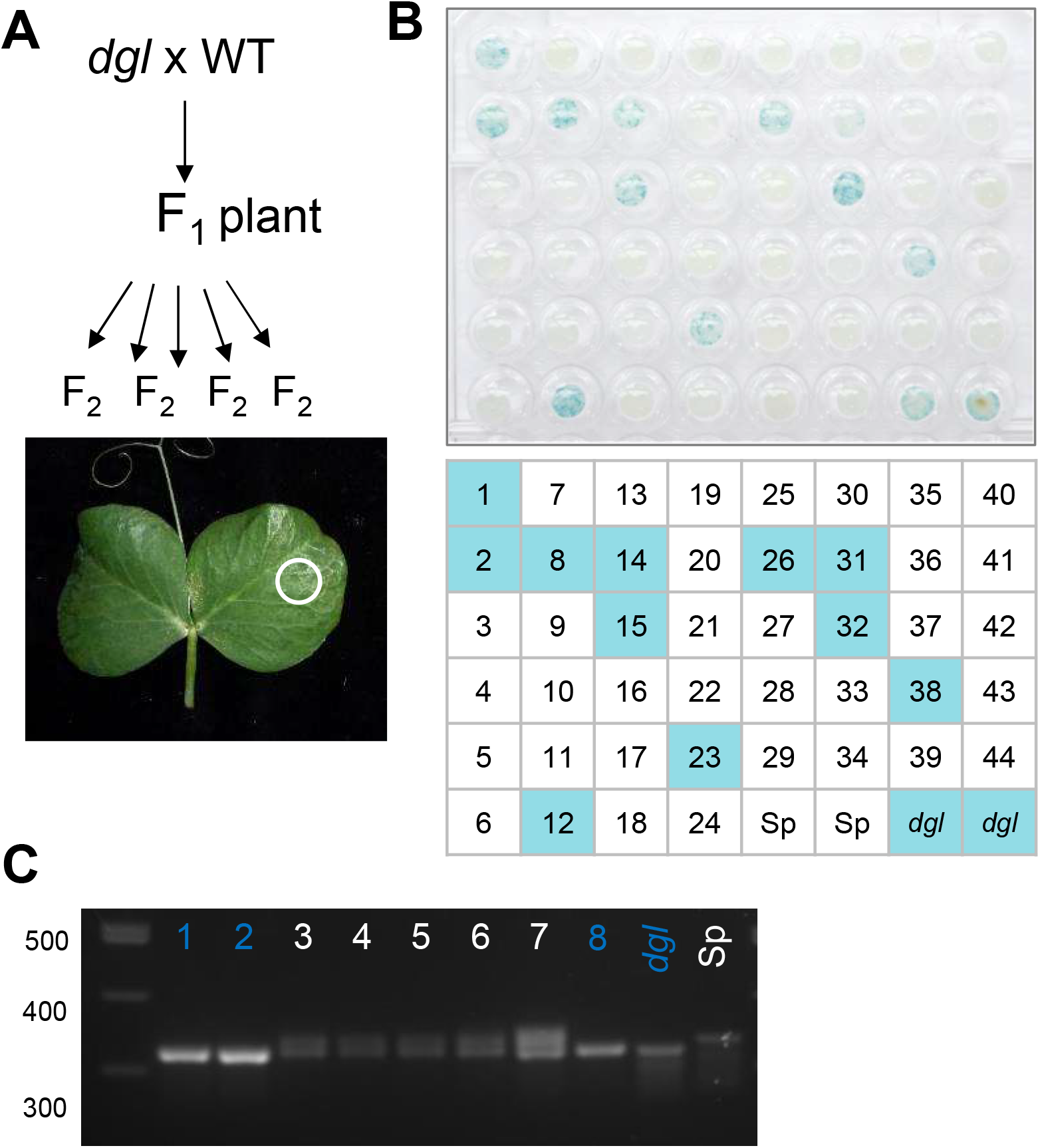
Co-segregation analysis of the *dgl* mutation and iron accumulation in pea, *Pisum sativum* L. A. The *dgl* mutant was crossed with a pea variety acting as wild-type for the locus (JI804), to obtain an F2 population of 44 plants. B. Discs (3 mm diameter) of the second leaf were stained for iron and scored for the iron-accumulating phenotype. C. PCR analysis of selected F_2_ plants, *dgl* and the wild-type control Sparkle (Sp), to detect the 15-bp deletion in *Psat1g036240* as well as the wild-type allele.

**Supplemental Figure 3.**
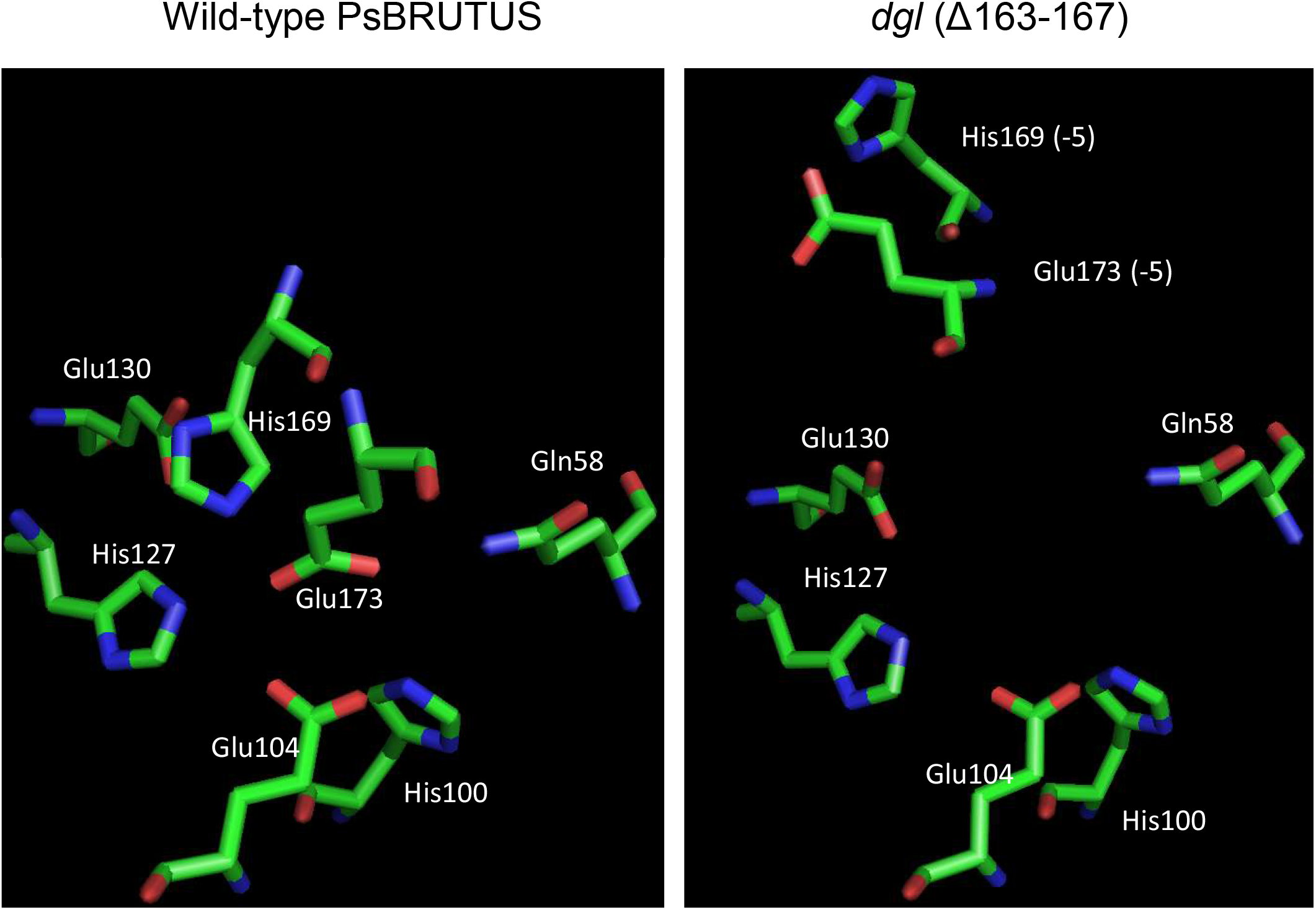
Models to show the position of amino acid ligands of the diiron centre in the hemerythrin 1 domain, in wild-type pea BRUTUS (left) and as a consequence of the 5-aa deletion in *dgl* (right). The diiron centre is predicted to have 7 ligands (plus water or oxygen) following a pattern that is conserved in all hemerythrins (H…HxxxE…H…HxxxE), consisting of histidine (His, H), glutamate (Glu, E) plus one glutamine (Gln). Because of the 5 amino acid deletion in the *dgl* mutant, the nearby His169 and Glu173 are predicted to be displaced and pointing away from the active site.

**Supplemental Figure 4.**
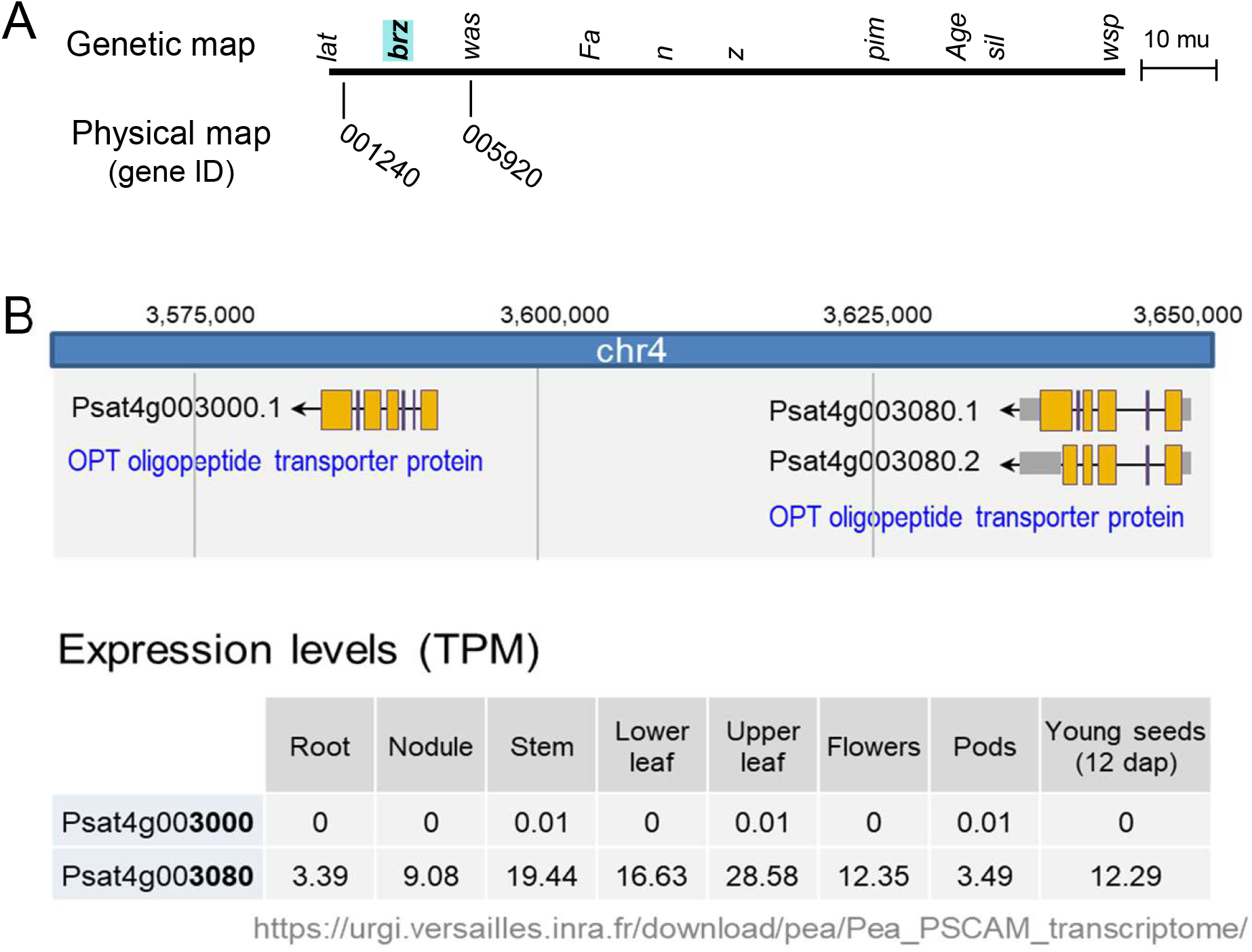
The pea homolog *OPT3* is a candidate gene for *BRZ*. A. The *brz* mutation was previously mapped to the tip of chromosome 4, between the genetic markers *lat* and *was* (Kneen et al., 1990; Ellis & Poyser, 2002). B. Detail of chromosome 4 showing the two neighboring *OPT3* paralogs, *Psat1g003000* and *Psat1g003080*. Expression data in transcripts per million (TPM) from https://urgi.versailles.inra.fr/ indicate that only one of the two paralogs, *Psat1g003080*, is expressed.

**Supplemental Figure 5.**
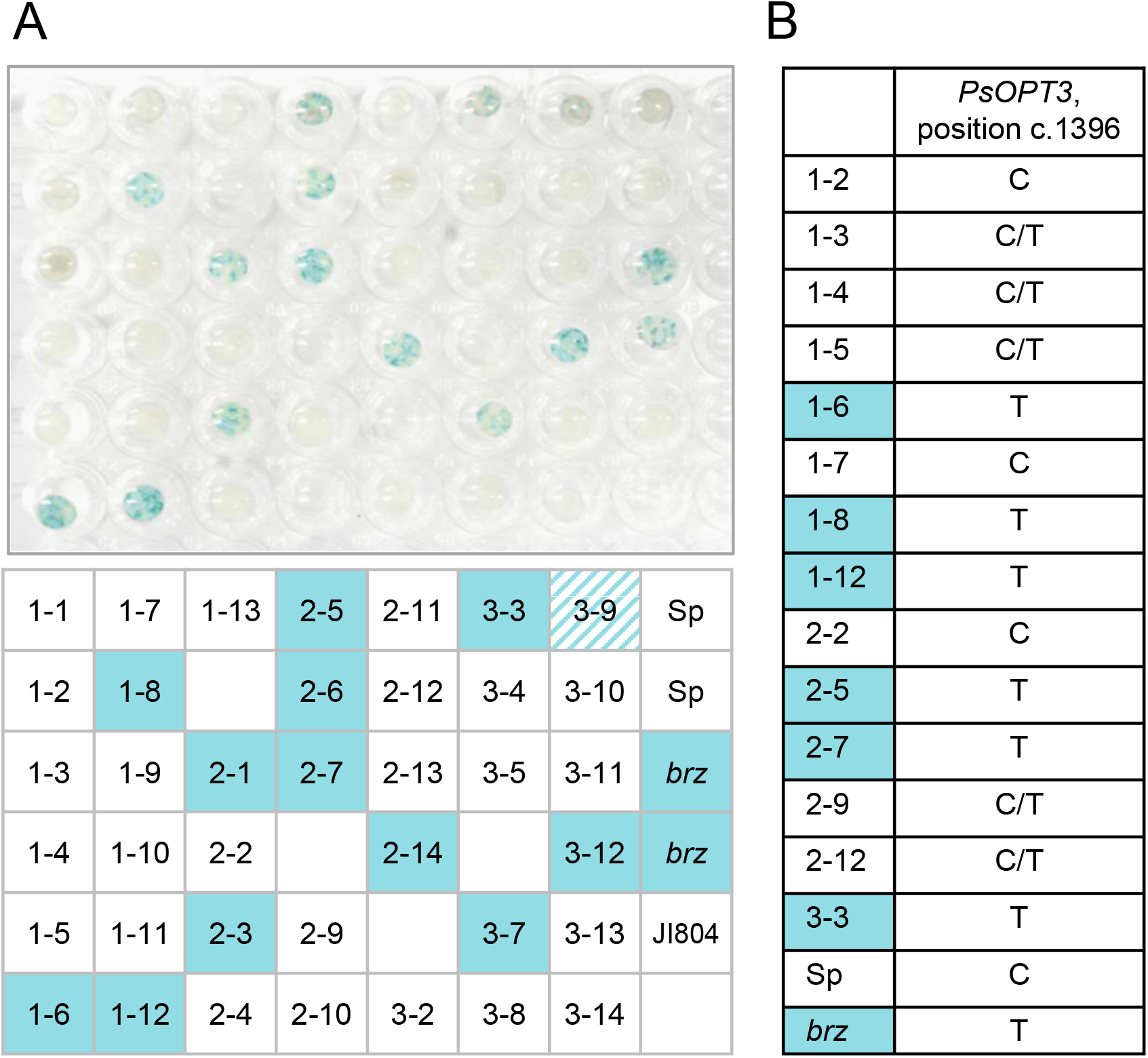
Co-segregation analysis of the *brz* mutation and iron accumulation in pea, *Pisum sativum* L. A. Iron-stained leaf discs (3 mm diameter) of F_2_ plants from crosses between *brz* and JI804 used as wild type. The F_2_ were from 3 different F1 plants. B. Allele variation for c.1396 in *Psat4g003080* / *PsOPT3* in 14 F_2_ plants, wild-type Sparkle (Sp) and *brz* serving as negative and positive controls, respectively.

**Supplemental Figure 6.**
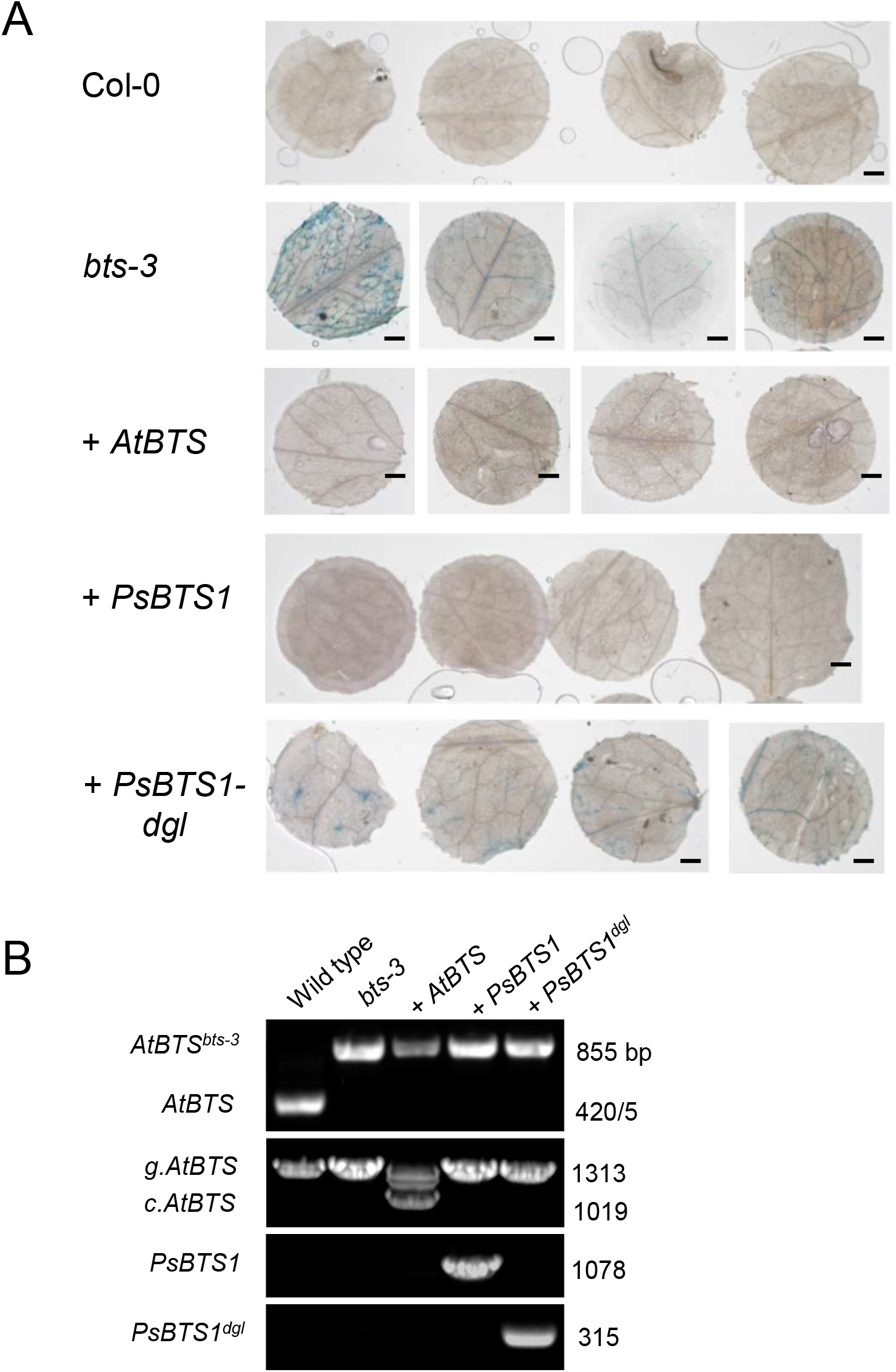
Genetic complementation of the Arabidopsis *bts-3* mutant with pea *BTS1*. **A**. Heterozygous Arabidopsis *bts-3* plants were transformed with T-DNAs containing cDNA sequences of Arabidopsis *BTS* (*AtBTS*), pea (*Ps*) *BTS1* and the *dgl-*associated variant of pea *BTS1* (*PsBTS1-dgl*). T1 plants homozygous for *bts-3* were selected, and T2 seedlings from these plants were grown up. Leaf discs (3 mm) of older leaves were stained for iron and imaged by light microscopy. Representative images of 2 independent lines are shown. Scale bar is 0.5 mm. **B**. Genotyping of the plants pictured in Fig. 4, using PCR. The *bts-3* mutation removes a PflMI restriction site, detected by PCR followed by PflMI treatment. Sizes of the nucleotide bands are on the right. Primers are listed in Suppl. Table 3.

**Supplemental Figure 7.**
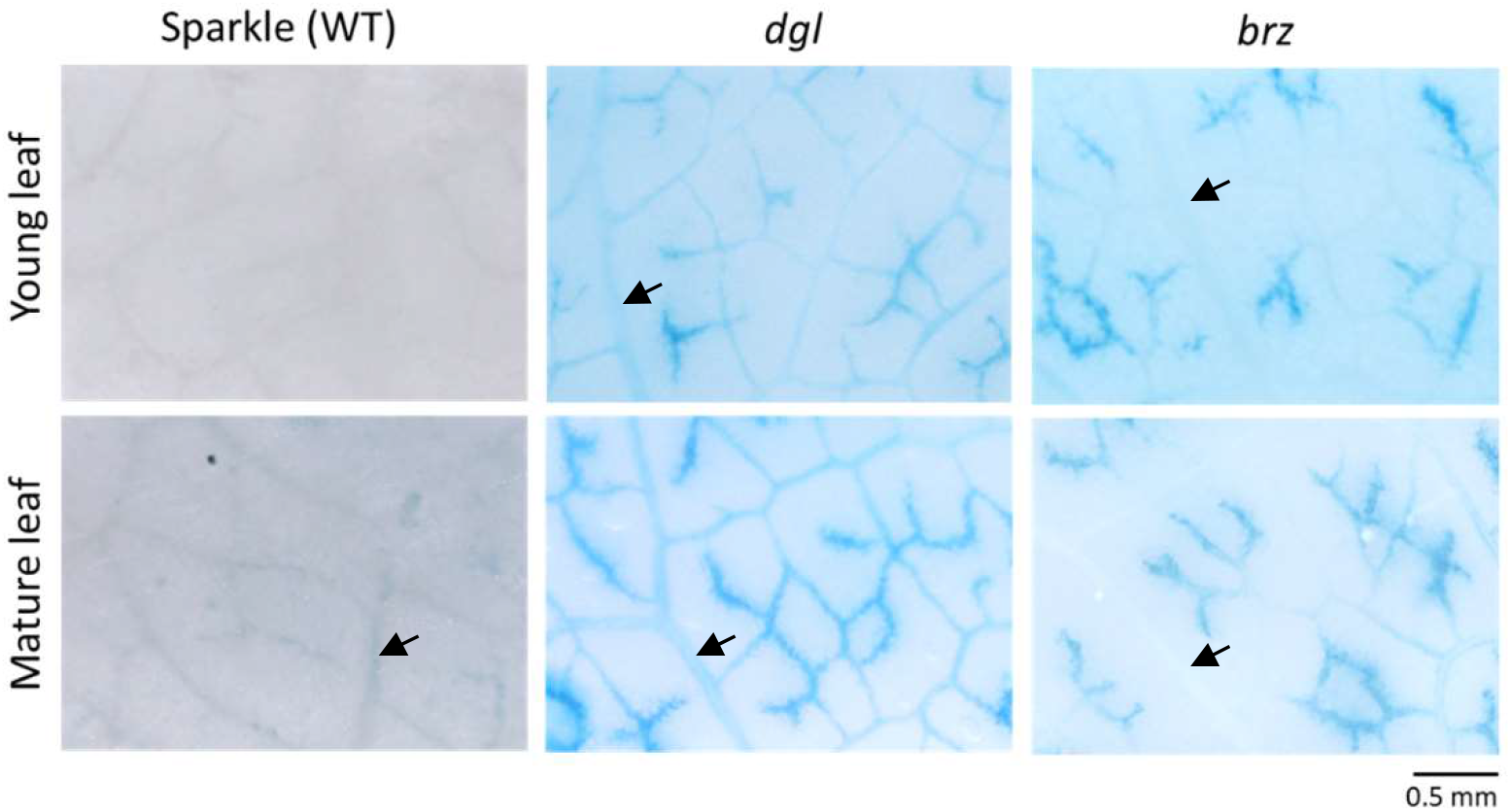
Pattern of iron accumulation in the leaves of *dgl* and *brz* mutants. Leaves were stained using the Perls’ method and imaged using a stereo microscope and bright-field illumination. Larger veins (arrows) accumulate iron in the *dgl* mutant but not in the *brz* mutant.

**Supplemental Table 1.** Gene Ontology terms of differentially expressed genes in leaves from the pea dgl mutant compared to the corresponding wild-type variety Sparkle. BP, biochemical pathway; MF, molecular function.

| category | term | ontology | over_rep_padj |
| --- | --- | --- | --- |
| GO:0006826 | iron ion transport | BP | 3.58E-07 |
| GO:0006879 | cellular iron ion homeostasis | BP | 4.95E-06 |
| GO:0008199 | ferric iron binding | MF | 4.95E-06 |
| GO:0055072 | iron ion homeostasis | BP | 4.95E-06 |
| GO:0000041 | transition metal ion transport | BP | 1.28E-05 |
| GO:0030003 | cellular cation homeostasis | BP | 2.25E-05 |
| GO:0055082 | cellular chemical homeostasis | BP | 2.25E-05 |
| GO:0006873 | cellular ion homeostasis | BP | 2.64E-05 |
| GO:0050801 | ion homeostasis | BP | 8.73E-05 |
| GO:0055080 | cation homeostasis | BP | 8.73E-05 |
| GO:0048878 | chemical homeostasis | BP | 0.000207 |
| GO:0030001 | metal ion transport | BP | 0.005288 |

**Supplemental Table 2.**
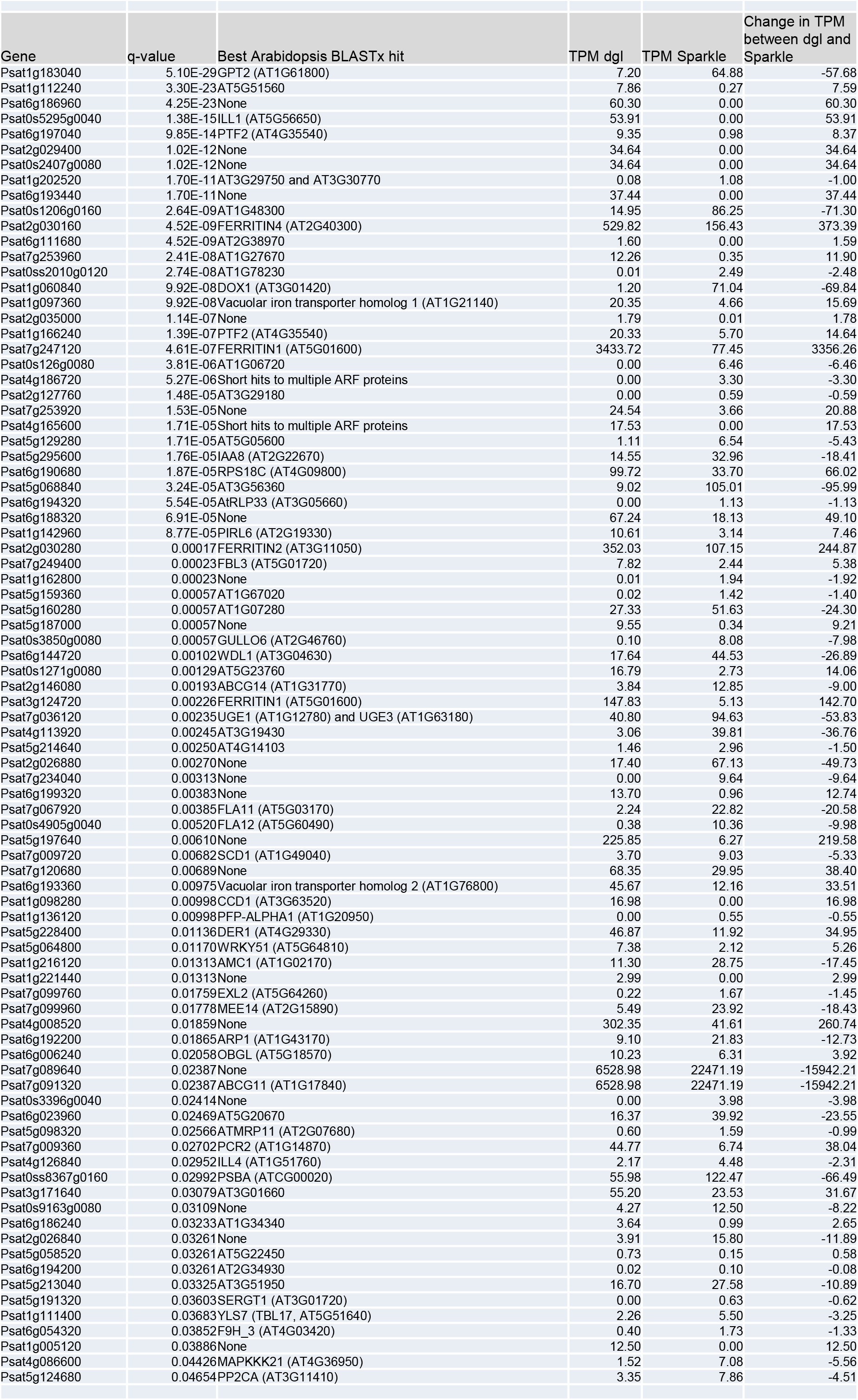
Differentially expressed genes in leaves from the pea dgl mutant compared to the corresponding wild-type variety Sparkle.

**Supplemental Table 3.**
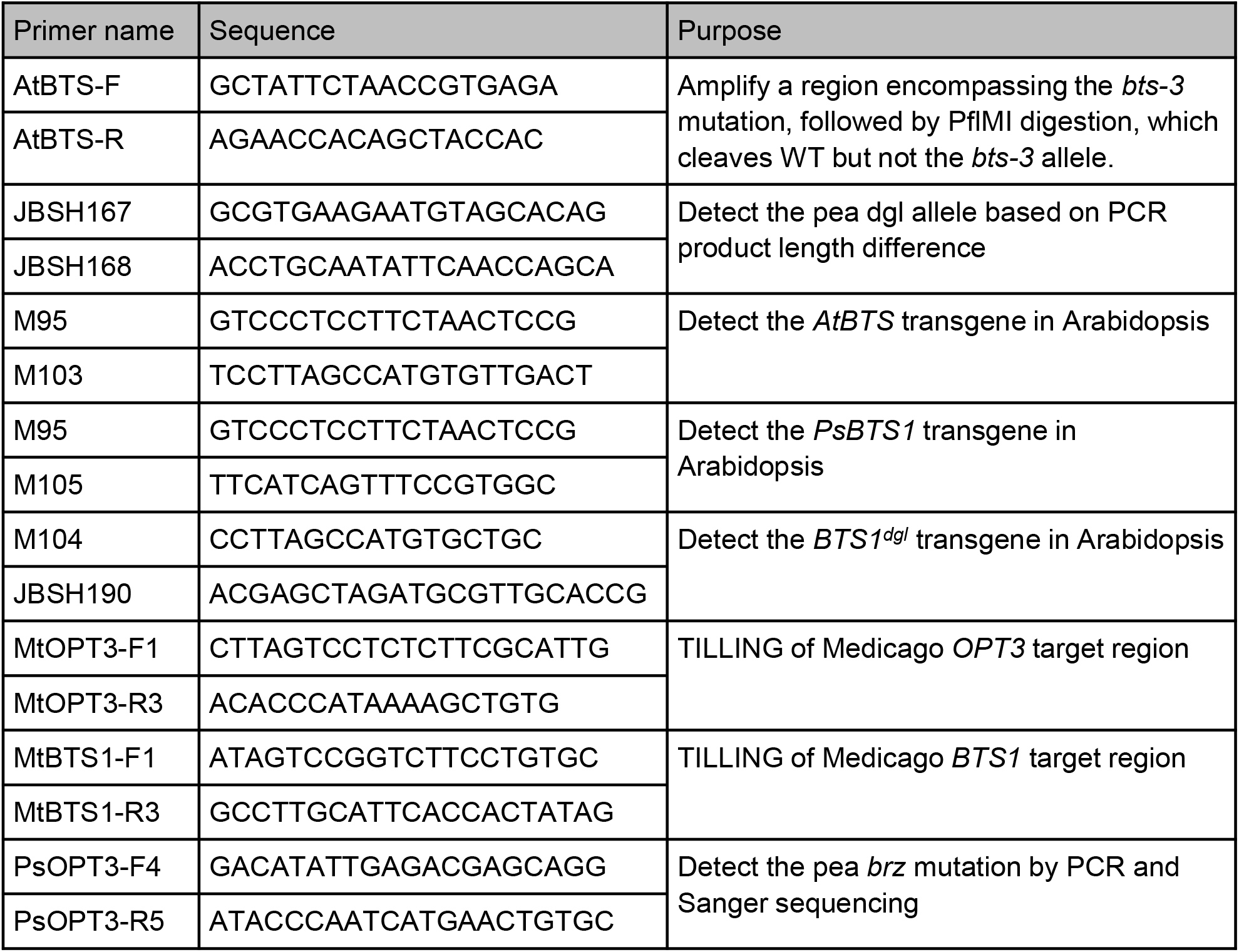
Primers used in this study.

**Supplemental Table 4.** TILLING mutations in *Medicago truncatula* genes.

| Medicago *BTS1* / *Medtr6g083900* | Amino acid change | Phenotype in Medicago |
| --- | --- | --- |
| c.G2188A | p.Asp730Asn | Embryo lethal |
| c.G2419A | p.Gly807Arg | None* |
| c.G2734A | p.Asp912Asn | None |
| c.G2800A | p.Leu934Phe | None |
| Medicago *OPT3* / *Medtr4g133968* |  |  |
| c.G1219A | p.Asp407Asn | None |
| c.C1586T | p.Pro529Leu | Iron accumulation, bronze spots on older leaves, impaired growth |

